# Rapid Assessment of Target-Binding Fractions in Theranostic and Imaging Agents Using Size-Exclusion HPLC

**DOI:** 10.64898/2026.01.23.699790

**Authors:** Ashtyn McAdoo, Kamal Jouad, Eben L. Rosenthal, Adam J. Rosenberg

**Affiliations:** Chemical and Physical Biology, Vanderbilt University, Nashville, TN USA; Department of Otolaryngology-Head and Neck Surgery, Vanderbilt University Medical Center, Nashville, TN, USA; Vanderbilt University Institute of Imaging Science, Vanderbilt University Medical Center, Nashville, TN USA; Department of Radiology and Radiological Sciences, Vanderbilt University Medical Center, Nashville, TN, USA

**Author notes:** Corresponding Author: Adam J. Rosenberg, Vanderbilt University Medical Center, Nashville TN, USA.

**Keywords:** Immunoreactivity, Target-binding fraction, Lindmo assay, Quality Control, Radiochemistry, Radioimmunoconjugates

## Abstract

**Background:** The clinical translation of molecularly targeted therapeutics and imaging agents represents a cornerstone of precision oncology, with the global theranostics market projected to exceed $25 billion by 2030. However, the development of theragnostic agents or diagnostic companions remains constrained by analytical bottlenecks in quality control, such as target-binding specificity, which are increasingly required by regulatory agencies as product release criteria during the translation process. Current methods, including enzyme-linked immunosorbent assay (ELISA), which require specialized resources or external CROs, or bead-based assays for radiolabeled compounds, which involve complex multi-step protocols; these limitations and others hamper their practical implementation in clinical manufacturing environments. Assay delays can postpone clinical trial initiation, increase development costs, and delay patient access to these agents.

**Results:** We have developed and validated a rapid, size-exclusion high-performance liquid chromatography (SE-HPLC) method for the determination of target-binding fractions of labeled biologics. The method separates the unbound biologic from the larger antigen-bound complex, allowing for rapid quantification. We validated the method using a panel of fluorescently labeled antibodies (panitumumab-IRDye800CW, nivolumab-IRDye800CW) and radiolabeled biologics ([18F]GEH200521, [18F]NOTA-ABY-030), assessing linearity, specificity, and concentration independence.

The SE-HPLC method achieved excellent separation of bound and unbound species with a resolution (Rs) of 3.2. A strong linear relationship (R^2^ = 0.999) was observed between the antigen-to-antibody ratio and the measured binding fraction. The method demonstrated high specificity, with no binding detected with non-target antigens. The total assay and analysis time was less than 35 minutes, a significant improvement over traditional methods.

**Conclusions:** SE-HPLC provides a rapid, specific, and cost-effective alternative to traditional binding fraction assessment methods, reducing quality control timelines from weeks/hours to minutes. The method’s compatibility with both fluorescent and radiolabeled biologics and integration with existing HPLC infrastructure represents a significant advancement in development workflows.

## Background

The era of precision oncology has been defined by the development of molecularly targeted therapeutics, which have transformed cancer treatment paradigms.[1-12] The global market for these agents, including antibody-drug conjugates, immune checkpoint inhibitors, and radiolabeled theranostics, is projected to exceed $100 billion in 2024.[13] The clinical success of these biologics is contingent upon their ability to bind to their intended molecular targets with high specificity and affinity. Therefore, a critical component of the manufacturing process is the analytical measure of the target-binding fraction / potency, a key quality attribute that is mandated by regulatory agencies.[14] The clinical success of targeted therapeutics is dependent on the interaction between the targeting vector and its intended target. These therapeutics require analytical validation to ensure that the targeting vector maintains its binding specificity throughout the manufacturing process. When these biologics are conjugated with imaging or therapeutic labels, additional analytical validation is required to ensure the labeling process does not alter target-binding integrity. The binding fraction (the percentage of labeled biologic that retains the ability to bind its intended target fraction) serves as a critical quality measure that correlates with therapeutic efficacy and patient safety. The regulatory framework for targeted biologics mandates rigorous assessment of binding specificity for patient safety and therapeutic efficacy[14]. The United States Food and Drug Administration (FDA), European Medicines Agency (EMA), and other regulatory authorities require reproducible analytical validation demonstrating that labeled biologics maintain their target-binding properties throughout the manufacturing process and product lifecycle.[15-17]

To assess the functional integrity of the labeled biologic, reactivity assays like ELISAs involve multiple incubation, blocking, and washing steps that are labor-intensive and time-consuming.[18] The Lindmo assay, widely used for radiolabeled compounds, relies on cell-based systems requiring specialized cell culture facilities, trained personnel, and extensive preparation time.[19-21] More recently developed bead-based assays, while offering some improvements in throughput, still involve multi-step protocols and specialized equipment that limit their practical implementation.[22]

High-performance liquid chromatography (HPLC) represents a mature, widely available analytical platform that is already integral to the quality control of biologics manufacturing.[23, 24] When a targeting biologic binds to its intended antigen, the resulting complex has a substantially larger molecular weight than the unbound biologic, enabling chromatographic separation.[25, 26] Previously, researchers at Genentech developed an HPLC-based assay to measure the target-binding fraction of a zirconium-89-labeled minibody.[26] Here we describe the development, validation, and clinical implementation of a size-exclusion HPLC method for rapid assessment of target-binding fractions in both fluorescently labeled and radiolabeled biologics.

## Results

### Chromatographic Method Optimization and Column Performance Evaluation

To establish an HPLC-based quantification of the target-binding fraction, we evaluated two size-exclusion chromatography (SEC) columns. Initial assessment using a Biozen 1.8 μm dSEC-2 column achieved adequate separation of panitumumab-IRDye800CW from its EGFR complex, with retention times of 7.2 ± 0.1 minutes for the unbound antibody and 5.7 ± 0.1 minutes for the antibody-antigen complex, respectively. While the peak symmetry was excellent (tailing factors < 1.2), the resolution (Rs) was 1.8, which provided a limited margin for method robustness.

In contrast, the Superdex 200 Increase 10/300 GL column demonstrated superior separation performance. Using this column, the unbound panitumumab-IRDye800CW eluted at 14.1 ± 0.1 minutes, while the antibody-EGFR complex eluted at 10.4 ± 0.1 minutes, yielding an improved resolution of 3.2 (Figures 1, S1, and S2). Based on this enhanced performance, the Superdex 200 column was selected for further evaluation.

**Figure 1:**
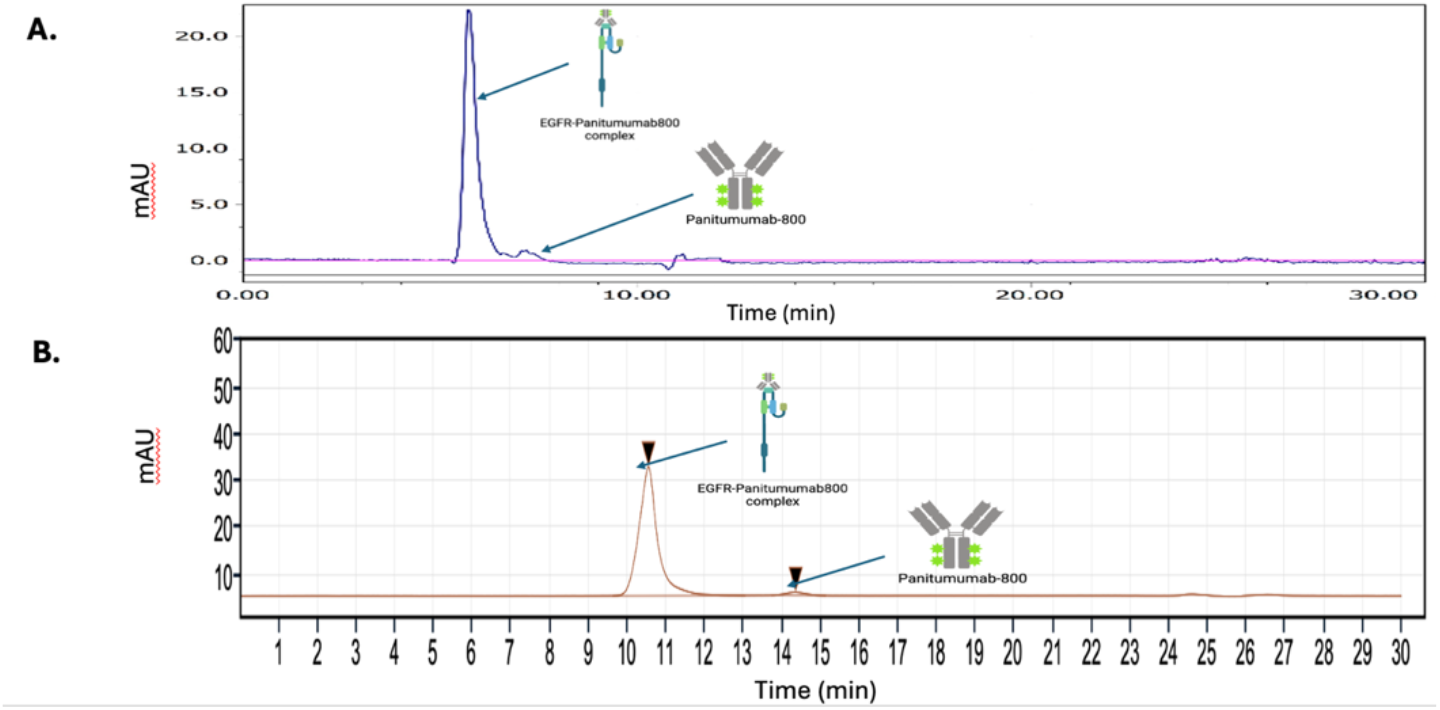
Representative chromatograms of panitumumab-IRDye800CW incubated with and without EGFR, showing the separation of the antibody-antigen complex from the unbound antibody on theA) Biozen 1.8 μm dSEC-2 and B) Superdex 200 column. The high resolution (Rs = 3.2) allows for accurate quantification of both species.

### ELISA Validation

To validate the results for the SE-HPLC method, the binding affinity by ELISA for pan800 was measured. A sample was shipped to an established CRO (RayBioTech), where EGFR binding was evaluated using an established ELISA protocol (successful in prior IND applications). The Pan800 met its binding affinity requirements, similar to unconjugated panitumumab, and demonstrated that the lot of Pan800 was suitable to be used as a standard. The results demonstrated that the binding affinity of Pan800 to EGFR was comparable to that of the unconjugated panitumumab, confirming its suitability as a 100% binding standard for the development of the SE-HPLC assay.

### Antigen to Antibody Ratio

The linearity of the SE-HPLC method was assessed by analyzing samples with varying molar ratios of EGFR to panitumumab-IRDye800CW. While the antigen volume remained constant, increasing amounts of pan800 were added to the antigen solution and analyzed by HPLC to generate the antigen-to-antibody ratios tested. As shown in Figure 2, the percent area under the curve for both the bound (10.4 ± 0.1 minutes) and the unbound (14.1 ± 0.1 minutes) peaks varied proportionally to the input ratio, confirming method linearity.

**Figure 2.**
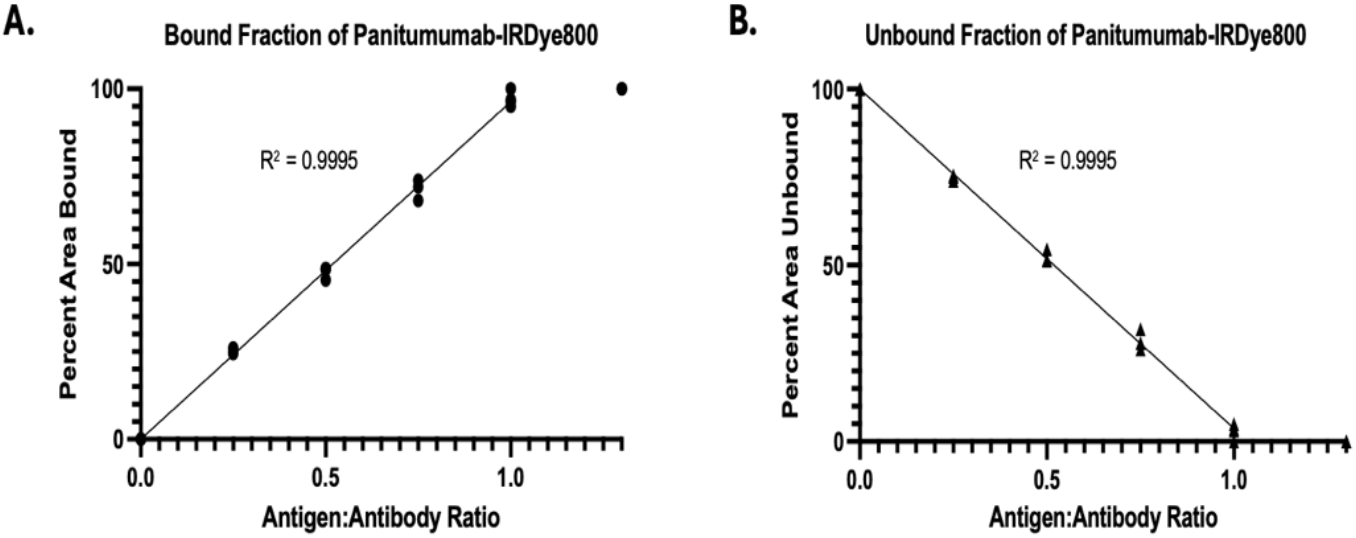
Linear regression analysis of the binding fraction of panitumumab-IRDye800CW as a function of the antigen-to-antibody ratio. The high correlation coefficient (R^2^ = 0.999) demonstrates the linearity of the method over the tested range.

A linear regression was performed to assess the relationship between the antigen-to-antibody ratio (between the ratios 0 and 1) and the percent area bound or the percent area unbound (representing the labeled antibody). (Table 1) The R^2^ value for each graph is 0.999, indicating the linearity of the relationship between the amount of antibody bound or unbound and the ratio of antigen-to-antibody. At a 1.3:1 ratio, where the antigen was in excess, the antibody was 100% bound, confirming the saturation of the system. All experiments were conducted in triplicate to ensure reproducibility, and the results were consistent across all replicates. Linear regression analysis of the data from the 0.25:1 to 1:1 ratios demonstrated an excellent linear relationship (R^2^ = 0.999, p < 0.001). The slope of the regression line was 94.2 ± 1.1%, which is close to the theoretical value of 100%, and the y-intercept of -1.8 ± 0.7% was not significantly different from zero (p = 0.052), indicating the absence of systematic bias in the method.

**Table 1:** The bound vs unbound peak areas at varying concentrations of panitumumab-IRDye800 using a 1:1 antigen-to-antibody ratio. Each concentration was run in triplicate.

| Panitumumab-IRDye800CW Concentration (mg/mL) | Percent Bound to Antigen (%) | Percent Unbound Antibody (%) |
| --- | --- | --- |
| 11.1 | 97.5 ± 0.8 | 2.5 ± 0.8 |
| 5.5 | 96.9 ± 3.1 | 6.2 ± 3.1 |
| 2.8 | 95.7 ± 0.7 | 4.3 ± 0.7 |
| 1.4 | 98.5 ± 1.5 | 1.5 ± 1.5 |
| 0.7 | 98.8 ± 0.7 | 1.2 ± 0.7 |
| Average | 97.4 ± 3.6 | 2.6 ± 3.6 |
| RSD | 3.7% |  |

#### Specificity

The specificity of the assay was confirmed through a series of negative control experiments (Table 2). When panitumumab-IRDye800CW was incubated with non-target antigens HER2 and PD-L1, no binding was detected, with 100% of the fluorescent signal remaining in the unbound fraction. Similarly, when EGFR was incubated with non-target antibodies, including the anti-amyloid antibody lecanemab-IRDye800CW and an isotype control antibody (IgG2-IRDye800CW), no binding was observed. (Table 2).

**Table 2:** Negative controls of both the antibody and antigen were used to ensure specificity of the antibody, panitumumab-IRDye800, to the antigen, EGFR.

| Antibody | Antigen | Percent Bound to Antigen (%) | Percent Unbound Antibody (%) |
| --- | --- | --- | --- |
| Panitumumab-IRDye800 | EGFR | 97.5 ± 0.8 | 2.5 ± 0.8 |
| IgG-IRDye800 | EGFR | 0 ± 0 | 100 ± 0 |
| Lecanumab-IRDye800 | EGFR | 0 ± 0 | 100 ± 0 |
| Panitumumab-IRDye800 | HER2 | 0 ± 0 | 100 ± 0 |
| Panitumumab-IRDye800 | PD-L1 | 0 ± 0 | 100 ± 0 |

#### Validation of the assay with additional fluorescent and radiolabeled biologics

To demonstrate the broad applicability of the SE-HPLC method, we tested it with a range of other fluorescent and radiolabeled biologics.

Nivolumab-IRDye800, an anti-PD-1 antibody, was incubated with its target antigen, PD-1. The resulting complex eluted at 12.2 ± 0.1 minutes, while the unbound antibody eluted at 14.3 ± 0.1 minutes, similar to the panitumumab system. (Figures S3 & S4) The analysis showed that 100% of the nivolumab-IRDye800 was bound to PD-1, demonstrating the utility of the method for other fluorescently labeled antibodies.

#### Validation in other targeted agents

The method was successfully applied to two different radiolabeled biologics. First, the immunoreactivity of [^18^F]GEH200521, a nanobody-based radiopharmaceutical targeting CD8, was assessed. Upon incubation with the CD8 antigen, the binding fraction was determined to be 97.55% ± 0.83%. (Figures S5 – S8) Second, the binding of [18F]NOTA-ABY-030, an affibody-based radiopharmaceutical targeting EGFR, was quantified. (Figures S9 & S10) The analysis showed a binding fraction of 89.67% (standard deviation = 1.46). These results confirm that the SE-HPLC method is compatible with radiolabeled biologics and can provide rapid QC data for these time-sensitive agents.

Purity analysis of [^18^F]GEH200521 showed a primary retention peak at 8.28 ± 0.17 minutes, representing 92.04% ± 4.14% of the sample. Upon incubation with CD8 antigen, the resulting [^18^F]GEH200521–CD8 complex exhibited a shifted retention time of 5.57 ± 0.04 minutes, while the unbound [^18^F]GEH200521 peak shifted slightly to 8.63 ± 0.13 minutes. The complex comprised 87.33% ± 4.14% of the total area, and the unbound antibody accounted for 3.72% ± 1.06 %. When considering only the peaks corresponding to the bound and unbound antibody, 97.55% ± 0.83% of [^18^F]GEH200521 was bound to CD8, with 2.45% ± 0.83% remaining unbound.

## Discussion

Because conventional methods of measuring binding specificity are costly and inefficient, we compared these methods to size-exclusion HPLC. We demonstrated this using both fluorescently labeled therapeutic antibodies (Pan800, Nivo800) and two radiolabeled theragnostic agents, such as [^18^F]GEH200521 and [^18^F]NOYA-ABY-030. Our findings demonstrate that this method provides a robust and reliable alternative to traditional binding assays, addressing a critical bottleneck in the quality control (QC) pipeline for the clinical translation of these agents. Our findings suggest this as an alternative to more time-intensive cell or bead-based assays for use in quality control (QC) assessments of clinical theranostic and imaging agents.

While the principle of using SE-HPLC to separate bound from unbound species based on molecular weight has been previously demonstrated to determine the immunoreactive fraction of ZED8, a ^89^Zr-labeled anti-CD8 antibody, for cGMP-compliant manufacturing.[26, 27] While the ZED8 study focused on a single radiolabeled antibody, we have extended SE-HPLC assay to fluorescently labeled antibodies (panitumumab-IRDye800CW and nivolumab-IRDye800), a radiolabeled nanobody ([^18^F]GEH200521), and a radiolabeled affibody ([^18^F]NOTA-ABY-030). Here we demonstrate the value of this technique with a range of scaffold sizes and types of labels.

The linearity of the relationship between the amount of antibody (bound and unbound fractions) and the antigen-to-antibody ratio had an R^2^ value of 0.999. A 1:1 molar ratio of EGFR to panitumumab-IRDye800 was adequate for assessing the binding fraction of panitumumab-IRDye800, although complete saturation was not achieved, potentially due to experimental variability of the Pan800 and/or concentration. Although variability in the concentration of reactants can also lead to inconsistent results, this was not observed in the current study. When the concentration of pan800 was varied between 0.7 mg/mL and 11.1 mg/mL, no significant differences were observed in the results, demonstrating consistency and reproducibility of the method across varying concentrations and productions of labeled panitumumab. This technique is especially relevant for radiolabeled biologics, where binding fraction detection is often performed at microgram to nanogram concentrations per milliliter. Furthermore, we had a total analysis time of under 35 minutes, which also benefits uses in radiolabeling, allowing a “same-day” release strategy.

Because the 280 nm channel detects all proteins, including both antibody and antigen, it is unsuitable for assessing binding specificity. In contrast, for optically labeled compounds, the 780 nm wavelength chromatogram enables selective detection of the labeled drug without interference from excess or unbound antigen. Likewise, the gamma detection channel offers comparable specificity for radiolabeled biologics, detecting only the radiolabeled compound. These channels allow for a more accurate evaluation of the labeled antibody’s capacity to bind its target antigen.

From an economic perspective, the SE-HPLC method offers some cost savings. As noted in our analysis, the per-sample cost of our method is approximately $75, a fraction of the cost of an outsourced ELISA ($1,200-$2,500) or even an in-house ELISA ($400-$600). By leveraging existing HPLC infrastructure, the capital investment required to implement this method is minimal. Furthermore, the combined speed and simplicity of the assay lowers the barrier to entry for academic labs and small biotechnology companies seeking to translate novel biologics into the clinic.

Our study is limited by performing the studies in a single laboratory, and an inter-laboratory comparison would be necessary to establish its reproducibility and facilitate broader adoption. Furthermore, while we validated our method against an ELISA standard, a direct, head-to-head comparison with a cell-based Lindmo assay for the same radiolabeled compound was not performed.

## Conclusion

Use of SE-HPLC for measurement of binding fraction is a simple, specific, and reliable cell-free approach for analyzing the binding interactions between labeled biologics and their respective antigens using existing equipment available in most cGMP-compliant labeling facilities. We demonstrate that this approach is compatible with both fluorescently and radiolabeled biologics, an advantage not offered by any other current assay, enabling streamlined quality control while maintaining compliance with current cGMP guidelines.

## Methods

### General

All chemicals were obtained from commercial suppliers and were of ACS or USP grade (Millipore-Sigma, USA). Size-exclusion chromatography columns were acquired from Cytiva, Waters, or Phenomenex and used according to the manufacturer’s specifications. Fluorescently labeled antibodies, including panitumumab-IRDye800CW and nivolumab-IRDye800CW, were prepared in-house. Radiolabeled biologics, including [89Zr]Panitumumab, [^18^F]GEH200521, and [^18^F]NOTA-ABY-030, were synthesized using in-house procedures in accordance with the appropriate INDs. Radiolabeling precursors and reference standards were sourced externally (GE Healthcare, Almac Sciences) or prepared in-house. Antigens were sources from commercial sources (Sino Biologic or R&D Systems) or provided by GE Healthcare.

### Mobile Phase Solution

A. Mobile phase A was prepared by mixing Potassium Phosphate (200 mM) and potassium chloride (250 mM) at a pH of 6.2. The solution was filtered through a 0.45 µm nylon membrane to remove any gases. The solution was transferred to the solvent reservoir. This solution was used as the mobile phase for the Biozen 1.8μm dSEC-2, 200 Å column defined below.
B. Mobile phase B was prepared by mixing sterile water, phosphate-buffered saline (0.05X PBS), sodium azide (10 mM), and sodium chloride (150 mM). The solution was filtered through a 0.45 µm nylon membrane to remove any gases. The solution was transferred to the solvent reservoir. This solution was used as the mobile phase for the Superdex Column defined below.

### Columns and HPLC

An Agilent 1260 Infinity II LC System or a Lablogic LOGI-Chrom system was used to collect data. The HPLC system was equipped with a diode array or multi-wavelength UV detector and a gamma detector. Two different columns were utilized to identify the best separation of the compounds.

A. A Biozen 1.8μm dSEC-2, 200 Å column (Phenomenex, Part No. 00H-4787-E0) composed of silica was used to test the separation of the panitumumab-IRDye800 peak from the panitumumab-IRDye800-EGFR complex. The column was primed with mobile phase A, described above. Then, 10 µL of the sample was injected onto the column. The flow rate was set to 0.35 mL/min at 25 °C.
B. A Superdex 200 Increase 10/300 GL (Cytiva, Part No. 28990944) was also used to test the separation of the antibody-antigen complex from the unbound antibody. The column was primed with mobile phase B described above, and 20 µL of the sample was injected onto the column. The flow rate was 0.8 mL/min at 25 °C.
C. A Waters Acquity Protein BEH SEC was used for [18F]GEH200521. The column was primed and eluted with a proprietary mobile phase and under proprietary conditions.

HPLC chromatograms were collected, with detection at 280 nM, and either 780 nM (optical) or gamma detection. The peaks of interest are identified and quantified using the area under each peak and assessed as a percentage of the total peak area on the 780 nm/gamma chromatograms.

#### Binding Assessment of Radiolabeled and Fluorescently Labeled Biologics

To assess the binding fraction of labeled biologics, both radiolabeled and fluorescently labeled compounds were incubated with excess antigen and analyzed by size-exclusion high-performance liquid chromatography (SE-HPLC). Analytes included radiolabeled agents such as [^18^F]GEH200521, a CD8-targeting nanobody, and [^18^F]NOYA-ABY-030, an EGFR-targeting affibody, as well as fluorescently labeled antibodies Panitumumab-IRDye800 (pan800), an anti-EGFR antibody, and Nivolumab-IRDye800 (nivo800), a PD-1-targeting antibody.

The antigens used include EGFR (Sino Biological Inc, Cat. #: 10001-H08H-UE), PD-1 (Bio-Techne R&D system, Cat #: 8986-PD-100), CD8 (provided by GEHC), PDL1 (Sino Biological Inc, Cat. #: 10084-H08H), and HER2 (R&D Systems, Cat. #: 1129-ER). The lyophilized antigens were reconstituted according to the manufacturers’ instructions.

For each assay, the labeled compound was combined with its corresponding antigen in a 1 mL Eppendorf tube using a pipette and vortexed for 3–5 seconds at 20-25 °C. The sample was then transferred into an HPLC vial and injected onto a primed size-exclusion column. Total sample preparation time was approximately 5 minutes, followed by a 20 or 30-minute HPLC run. Detection was performed at 280 nm, and gamma or fluorescence channels, as applicable, based on labeling modality. The overall assay design and methodology are depicted in Figure 1. The general timeline of the assays discussed in this paper is represented in Figure 2 (wash steps have been removed, and the simplest version of the assay is assumed).

## Supplementary Materials

The online version contains supplementary material available at XXXX

## Author Contributions

Conceptualization, A.J.R., E.R.; experimental design, A.J.R., K.J., A.M.; analysis, A.M., K.J.; validation, A.M., K.J.; writing – original draft preparation, A.M.; writing – review and editing, A.J.R., E.R., A.M., K.J.; funding acquisition. A.J.R., E.R. All authors have read and agreed to the published version of the manuscript.

## Acknowledgments

We thank GE Healthcare for providing the GEH200521 materials. We would also like to thank the VUIIS Radiochemistry Core staff for performing some of the experiments; and Affibody AB for their expertise and material support for NOTA-ABY-030.

## Availability of data and materials

All data generated or analyzed during this study are included in this published article [and its supplementary information files].

## Declarations

### Ethics approval and consent to participate

Not applicable.

### Consent for Publication

Not applicable.

## References

1. Min H-Y, Lee H-Y. Molecular targeted therapy for anticancer treatment. Experimental & Molecular Medicine. 2022;54:1670–94. doi:10.1038/s12276-022-00864-3.

2. Lheureux S, Denoyelle C, Ohashi PS, De Bono JS, Mottaghy FM. Molecularly targeted therapies in cancer: a guide for the nuclear medicine physician. European Journal of Nuclear Medicine and Molecular Imaging. 2017;44:41–54. doi:10.1007/s00259-017-3695-3.

3. Renfro LA, An M-W, Mandrekar SJ. Precision oncology: A new era of cancer clinical trials. Cancer Letters. 2017;387:121–6. doi:10.1016/j.canlet.2016.03.015.

4. Orlova A, Magnusson M, Eriksson TLJ, Nilsson M, Larsson B, Höidén-Guthenberg I, et al. Tumor Imaging Using a Picomolar Affinity HER2 Binding Affibody Molecule. Cancer Research. 2006;66:4339–48. doi:10.1158/0008-5472.Can-05-3521.

5. Löfblom J, Feldwisch J, Tolmachev V, Carlsson J, Ståhl S, Frejd FY. Affibody molecules: Engineered proteins for therapeutic, diagnostic and biotechnological applications. FEBS Letters. 2010;584:2670–80. doi:10.1016/j.febslet.2010.04.014.

6. Day KE, Beck LN, Deep NL, Kovar J, Zinn KR, Rosenthal EL. Fluorescently labeled therapeutic antibodies for detection of microscopic melanoma. The Laryngoscope. 2013;123:2681–9. doi:10.1002/lary.24102.

7. Martelli C, Lo Dico A, Diceglie C, Lucignani G, Ottobrini L. Optical imaging probes in oncology. Oncotarget. 2016;7.

8. Solomon M, Liu Y, Berezin MY, Achilefu S. Optical Imaging in Cancer Research: Basic Principles, Tumor Detection, and Therapeutic Monitoring. Medical Principles and Practice. 2011;20:397–415. doi:10.1159/000327655.

9. Kircher MF, Hricak H, Larson SM. Molecular imaging for personalized cancer care. Molecular Oncology. 2012;6:182–95. doi:10.1016/j.molonc.2012.02.005.

10. Giuliani S, Paraboschi I, McNair A, Smith M, Rankin KS, Elson DS, et al. Monoclonal Antibodies for Targeted Fluorescence-Guided Surgery: A Review of Applicability across Multiple Solid Tumors. Cancers. 2024;16:1045.

11. Stupp R, Mason WP, Bent MJvd, Weller M, Fisher B, Taphoorn MJB, et al. Radiotherapy plus Concomitant and Adjuvant Temozolomide for Glioblastoma. New England Journal of Medicine. 2005;352:987–96. doi:doi:10.1056/NEJMoa043330.

12. Brahmer JR, Drake CG, Wollner I, Powderly JD, Picus J, Sharfman WH, et al. Phase I Study of Single-Agent Anti–Programmed Death-1 (MDX-1106) in Refractory Solid Tumors: Safety, Clinical Activity, Pharmacodynamics, and Immunologic Correlates. J Clin Oncol. 2023;41:715–23. doi:10.1200/jco.22.02270.

13. Targeted Therapy Market Competitive Analysis, Advancing Precision Medicine for Cancer and Beyond. 2025.

14. Analytical Procedures and Methods Validation for Drugs and Biologics. In: Research CfDEa, Research CfBEa, editors.; 2015.

15. Points to Consider in the Manufacture and Testing of Monoclonal Antibody Products for Human Use. In: Research CfBEa, Research CfDEa, editors.; 1997.

16. Delaney S, Grimaldi C, Houghton JL, Zeglis BM. MIB Guides: Measuring the Immunoreactivity of Radioimmunoconjugates. Molecular Imaging and Biology. 2024;26:213–21. doi:10.1007/s11307-024-01898-x.

17. Rhodes BA, Buckelew JM, Pant KD, Hinkle GH. Quality control test for immunoreactivity of radiolabeled antibody. Biotechniques. 1990;8:70–5.

18. Hornbeck PV. Enzyme-Linked Immunosorbent Assays. Current Protocols in Immunology. 2015;110:2.1.–2.1.23. doi:10.1002/0471142735.im0201s110.

19. Lindmo T, Boven E, Cuttitta F, Fedorko J, Bunn PA. Determination of the immunoreactive function of radiolabeled monoclonal antibodies by linear extrapolation to binding at infinite antigen excess. Journal of Immunological Methods. 1984;72:77–89. doi:10.1016/0022-1759(84)90435-6.

20. Mattes MJ. Limitations of the Lindmo method in determining antibody immunoreactivity. International Journal of Cancer. 1995;61:286–8. doi:10.1002/ijc.2910610224.

21. Dux R, Kindler-Röhrborn A, Lennartz K, Rajewsky MF. Determination of immunoreactive fraction and kinetic parameters of a radiolabeled monoclonal antibody in the absence of antigen excess. Journal of Immunological Methods. 1991;144:175–83. doi:10.1016/0022-1759(91)90084-S.

22. Sharma SK, Lyashchenko SK, Park HA, Pillarsetty N, Roux Y, Wu J, et al. A rapid bead-based radioligand binding assay for the determination of target-binding fraction and quality control of radiopharmaceuticals. Nuclear Medicine and Biology. 2019;71:32–8. doi:10.1016/j.nucmedbio.2019.04.005.

23. Lam C, Patel P. Food, Drug, and Cosmetic Act. StatPearls. Treasure Island (FL): StatPearls Publishing Copyright © 2025, StatPearls Publishing LLC.;2025.

24. Fekete S, Beck A, Veuthey J-L, Guillarme D. Theory and practice of size exclusion chromatography for the analysis of protein aggregates. Journal of Pharmaceutical and Biomedical Analysis. 2014;101:161–73. doi:10.1016/j.jpba.2014.04.011.

25. Zamora PO, Sass K, Cardillo AS, Lambert CR, Budd P, Marek MJ, et al. Affinity thin-layer chromatography test of the immunoreactive fraction of radiolabeled antibodies. Biotechniques. 1994;16:306–11.

26. Gill H, Seipert R, Carroll VM, Gouasmat A, Yin J, Ogasawara A, et al. The Production, Quality Control, and Characterization of ZED8, a CD8-Specific 89Zr-Labeled Immuno-PET Clinical Imaging Agent. The AAPS Journal. 2020;22:22. doi:10.1208/s12248-019-0392-0.

27. Kist de Ruijter L, van de Donk PP, Hooiveld-Noeken JS, Giesen D, Elias SG, Lub-de Hooge MN, et al. Whole-body CD8+ T cell visualization before and during cancer immunotherapy: a phase 1/2 trial. Nature Medicine. 2022;28:2601–10. doi:10.1038/s41591-022-02084-8.

