## Supplemental Files for "Rapid Assessment of Target-Binding Fractions in Theranostic and Imaging Agents Using Size-Exclusion HPLC"

### Panitumumab-IRDye800 – EGFR Binding

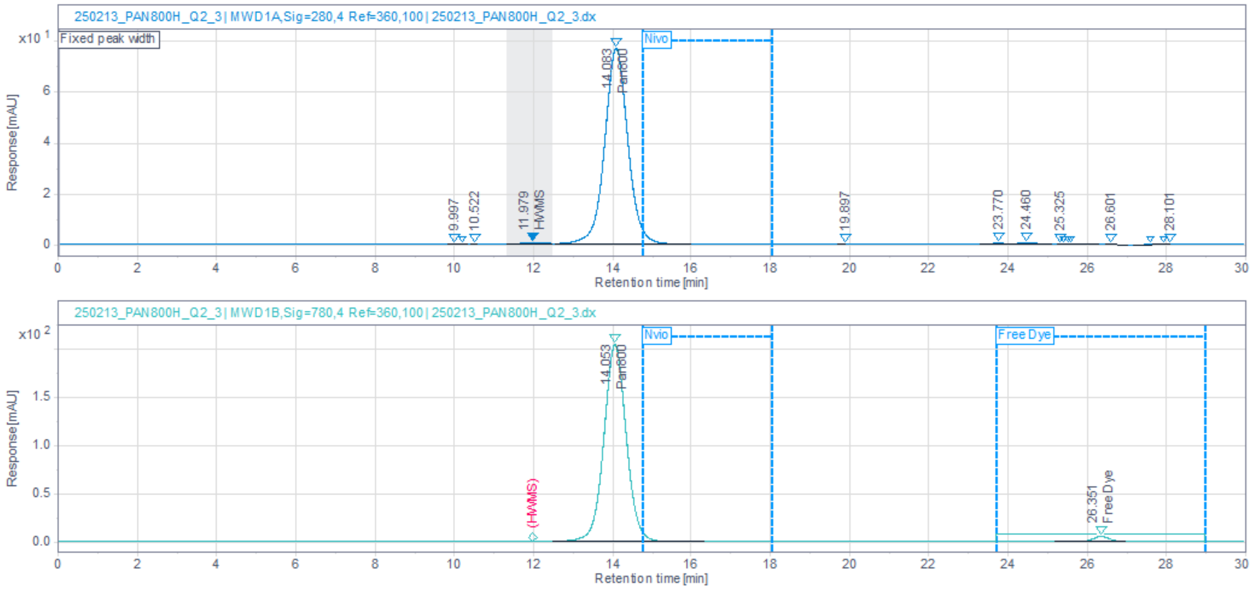


**Figure S1:** SEC of Panitumumab-IRDye800CW (Top, UV @ 280 nm; Bottom UV @ 780 nm)

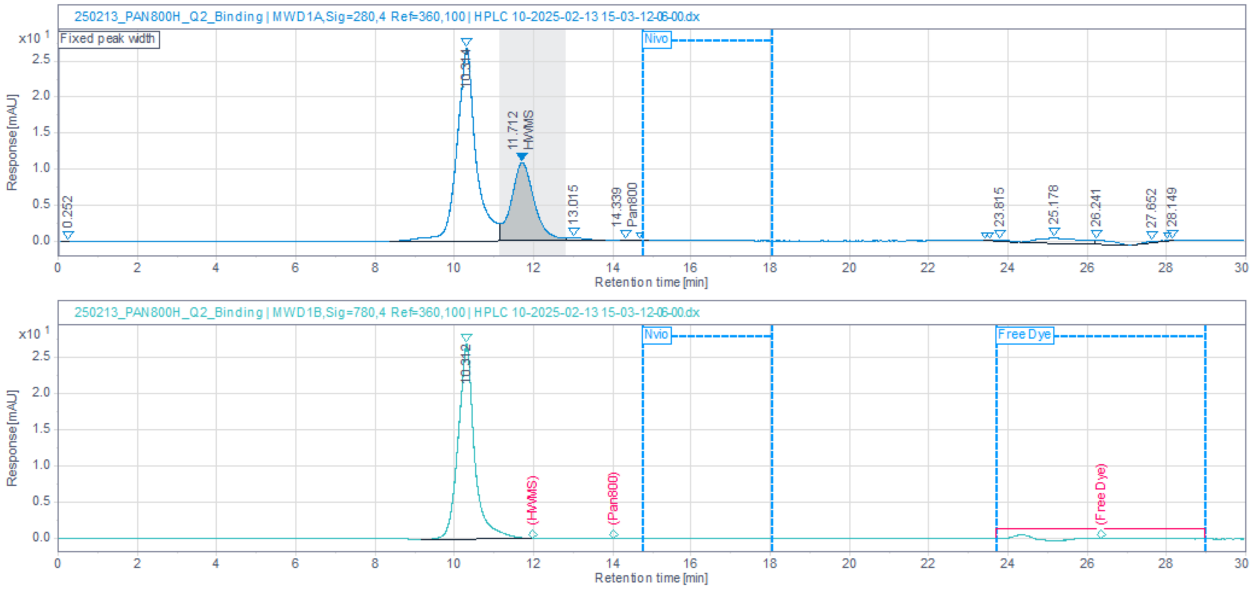


**Figure S2:** SEC of Panitumumab-IRDye800CW/EGFR Binding Assay (Top, UV @ 280 nm; Bottom UV @ 780 nm)

### Nivolumab-IRDye800 – PD-1 Binding

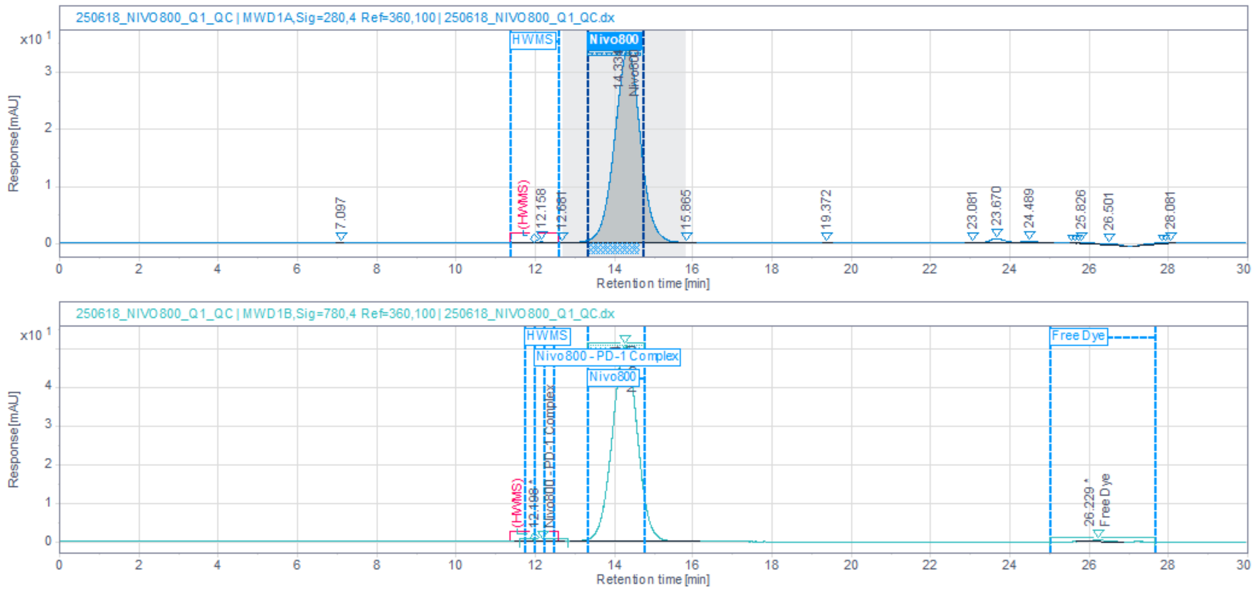


**Figure S3:** SEC of Nivolumab-IRDye800CW (Top, UV @ 280 nm; Bottom UV @ 780 nm)

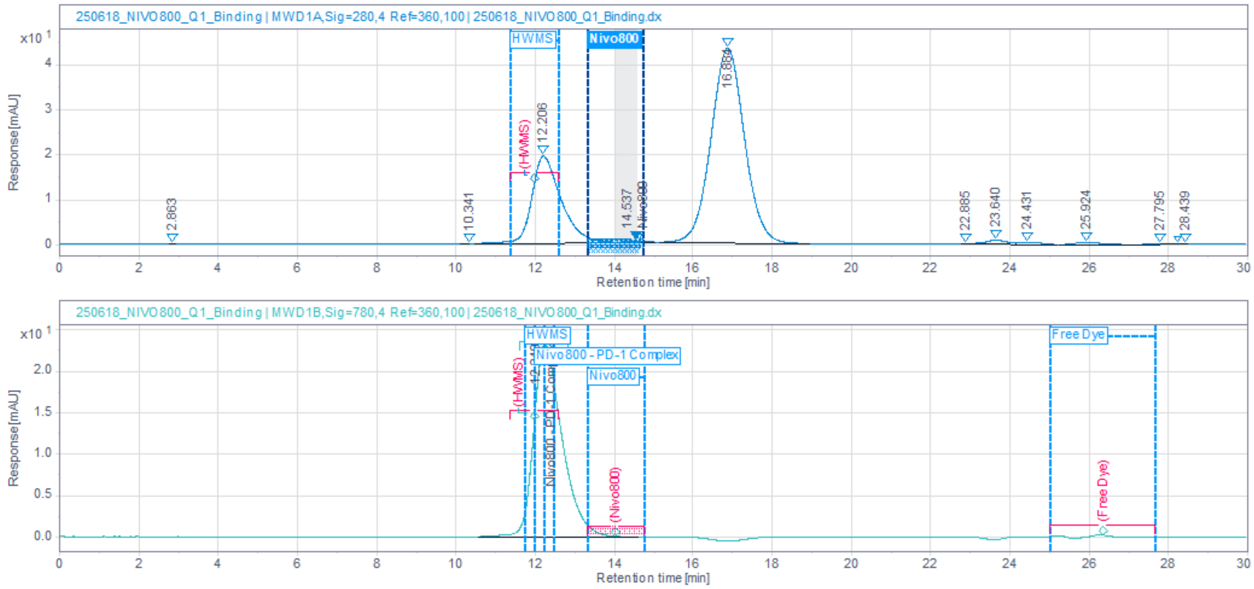


**Figure S4:** SEC of Nivolumab-IRDye800CW/PD-1 Binding Assay (Top, UV @ 280 nm; Bottom UV @ 780 nm)

**[18F]GEH200521 CD8 Binding**

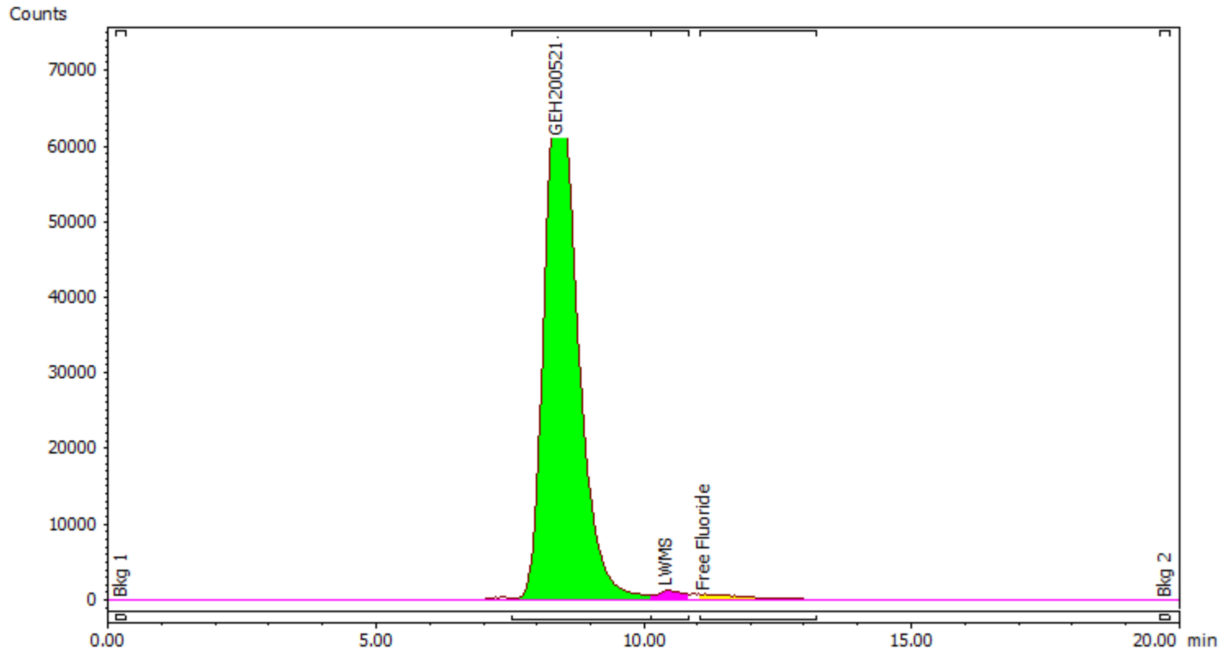


**Figure S5:** SEC of [18F]GEH200521 (Gamma Detector)

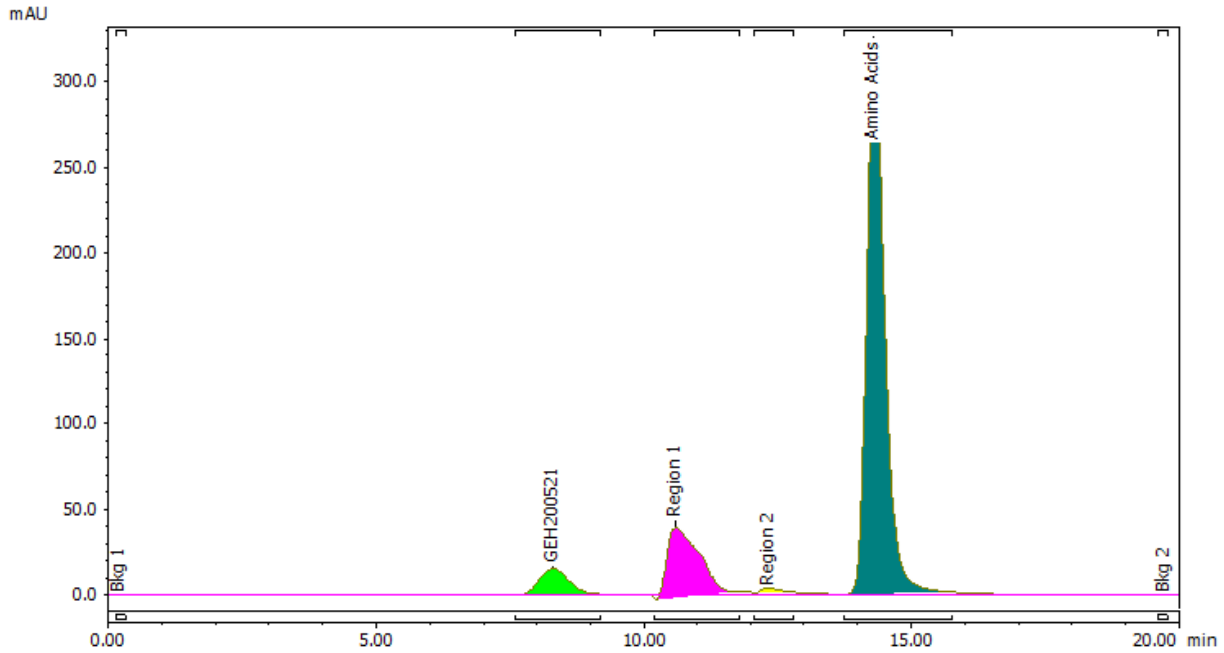


**Figure S6:** SEC of [18F]GEH200521 (UV @ 280 nm)

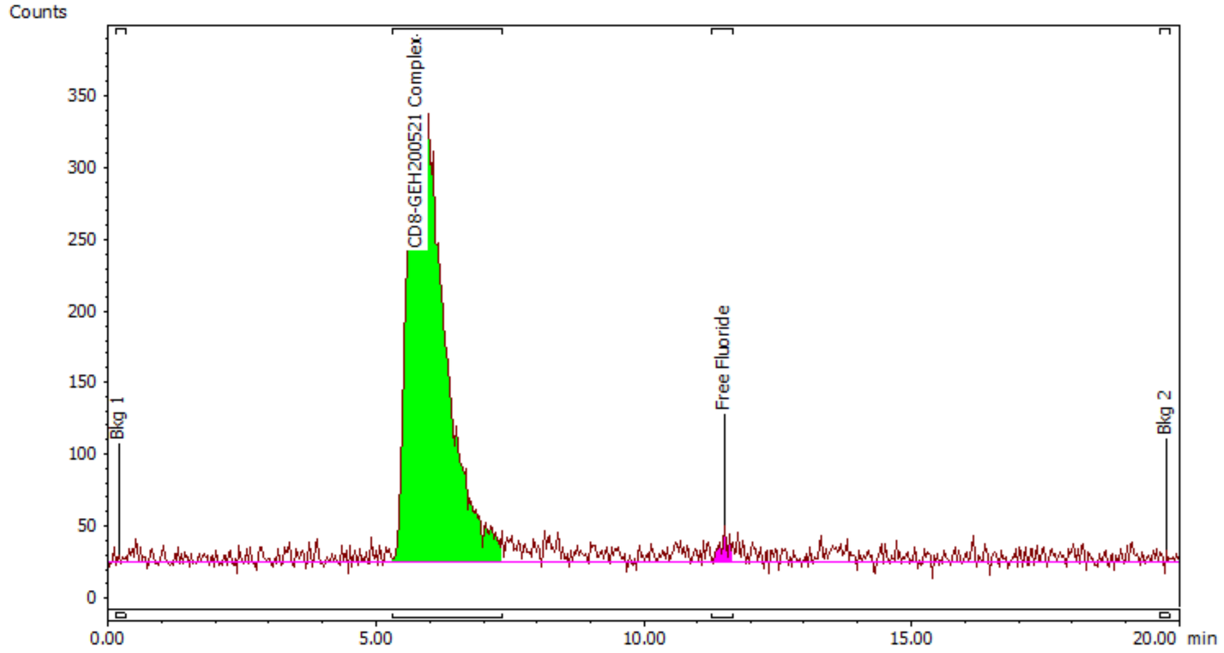


**Figure S7:** SEC of [18F]GEH200521/CD8 Complex (Gamma Detector)

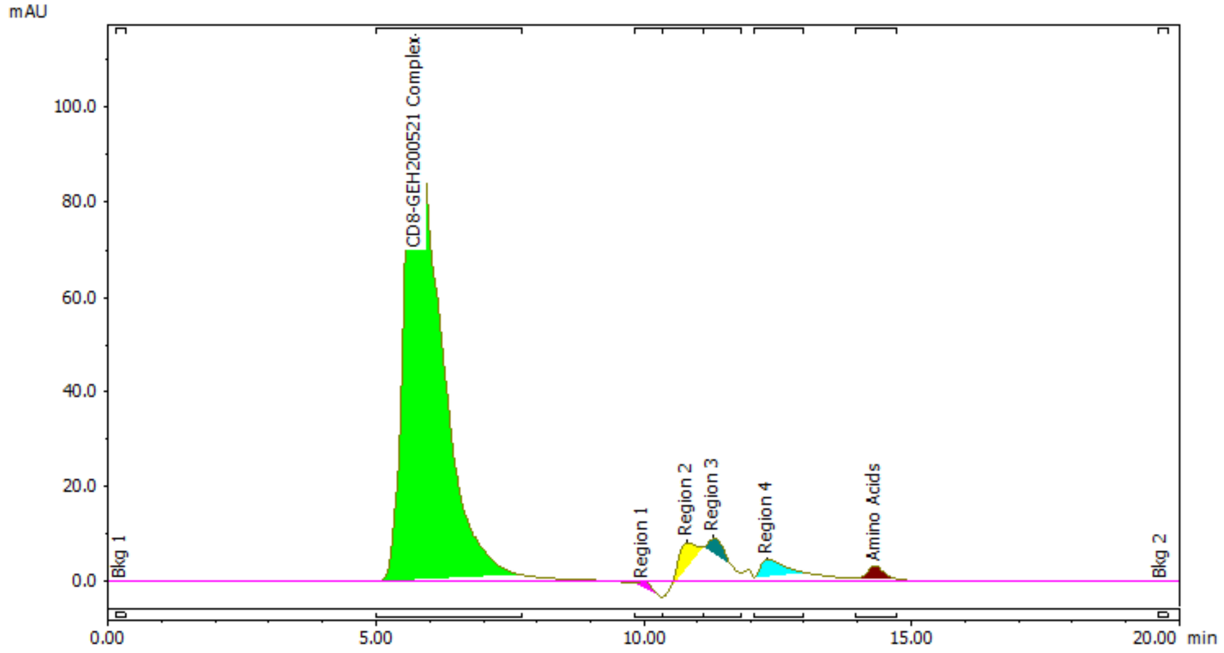


**Figure S8:** SEC of [18F]GEH200521/CD8 Complex (UV @ 280 nm)

**[18F]NOTA-ABY-030**

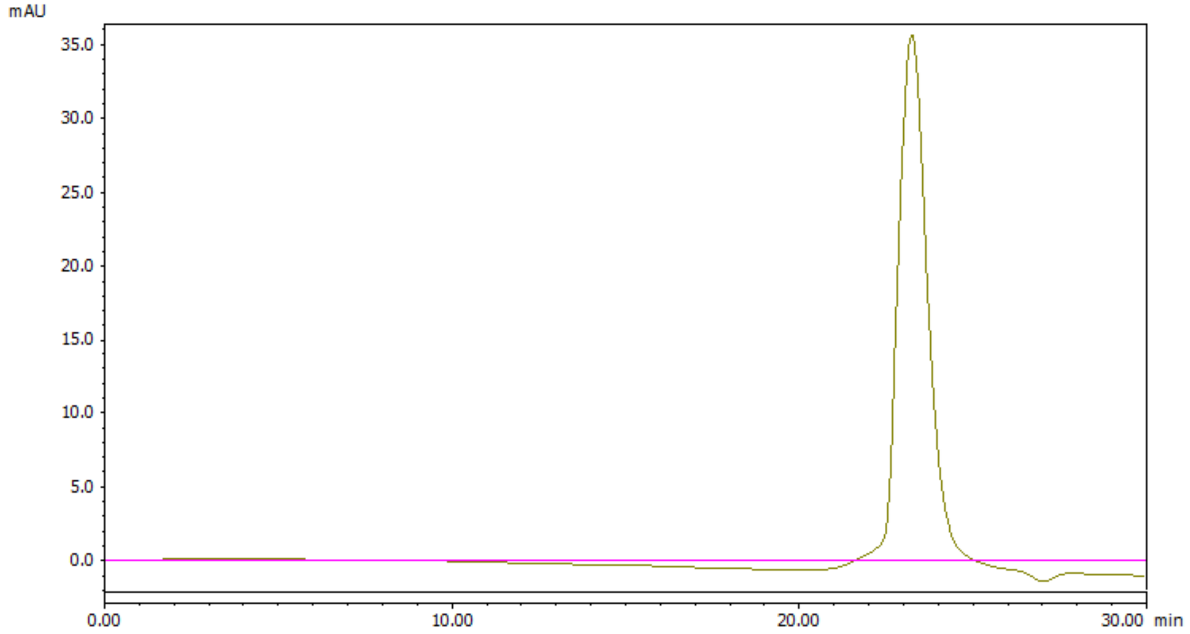


**Figure S9:** SEC of [18F]NOTA-ABY-030 (Gamma Detector)

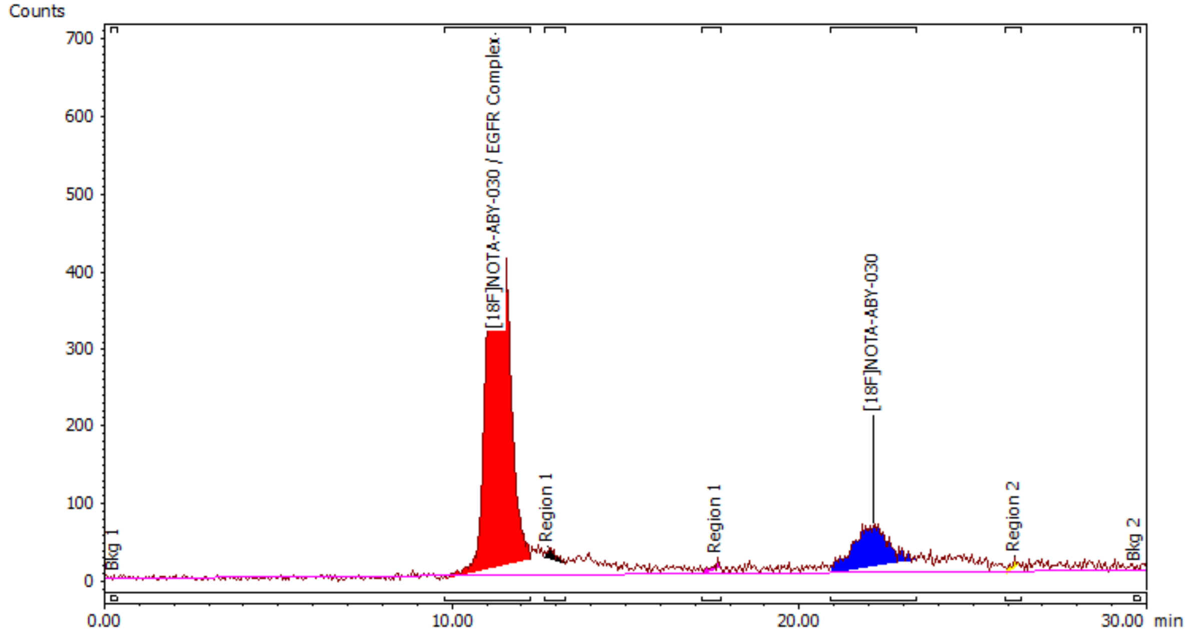


**Figure S10:** SEC of [18F]NOTA-ABY-030/EGFR Complex (Gamma Detector)
